# Generation and characterization of a novel mouse model of Becker Muscular Dystrophy with a deletion of exons 52 to 55

**DOI:** 10.1101/2023.11.16.567440

**Authors:** Lucie O. M. Perillat, Tatianna W. Y. Wong, Eleonora Maino, Abdalla Ahmed, Ori Scott, Elzbieta Hyatt, Paul Delgado-Olguin, Evgueni A. Ivakine, Ronald D. Cohn

**Author notes:** Equal contribution (Lucie O. M. Perillat, Tatianna W. Y. Wong). Corresponding authors: Ronald Cohn, Evgueni Ivakine. Co-senior authors: Evgueni Ivakine, Ronald Cohn ORCID: https://orcid.org/0000-0002-6775-1496.

## Abstract

Becker Muscular Dystrophy (BMD) is a rare X-linked recessive neuromuscular disorder caused by in-frame deletions in the *DMD* gene that result in the production of a truncated, yet functional, dystrophin protein. BMD is often considered a milder form of Duchenne Muscular Dystrophy, in which mutations typically result in the disruption of the reading frame and the malfunction or loss of dystrophin. The consequences of BMD-causing in-frame deletions on the organism are more difficult to predict, especially in regard to long-term prognosis. Here, we employed CRISPR-Cas9 technology to generate a new *Dmd* del52-55 mouse model by deleting exons 52-55, resulting in a typical BMD-like in-frame deletion. To delineate the long-term effects of this deletion, we studied these mice over 52 weeks. Our results suggest that a truncated dystrophin is sufficient to maintain wildtype-like muscle and heart functions in young mice. However, the truncated protein appears insufficient to maintain normal muscle homeostasis and protect against exercise-induced damage at 52 weeks. To further delineate the effects of the exons 52-55 in-frame deletion, we performed RNA-Seq pre– and post-exercise and identified several differentially expressed pathways that could explain the abnormal muscle phenotype observed at 52 weeks in the BMD model.

**Summary Statement:** We generated and characterized the long-term effects of a Becker Muscular Dystrophy-like in-frame deletion of exon 52 to 55 in mice.

## Introduction

Becker Muscular Dystrophy (BMD) is a rare X-linked recessive neuromuscular disorder caused by in-frame deletions in the *DMD* gene that result in the production of a truncated (or internally deleted), but functional, dystrophin protein. Dystrophin is an essential component of the dystrophin glycoprotein complex (DGC) that is located in the sarcolemma and required for muscle contraction and integrity (Fatehi et al., 2023). BMD is often considered as a milder form of Duchenne Muscular Dystrophy (DMD), in which mutations in the *DMD* gene, including large deletions and duplications, result in the disruption of the reading frame and the malfunction or loss of the dystrophin protein (Bushby et al., 1991). Young boys affected by DMD progressively lose their skeletal and cardiac muscular abilities, and require wheelchairs, ventilation and additional types of life support measures in their teenage years (Duan et al. 2021; Bello et al. 2016). While BMD tends to be milder than DMD, patients with BMD present with a wide spectrum of clinical presentations, ranging from asymptomatic to an early loss of ambulation (Bushby et al., 1991; Nigro et al., 1994). Levels of dystrophin expression vary between BMD patients and seem to correlate, to some extent, with disease severity (van der Bergen et al., 2014). This differential expression is also seen in individual patients, where levels of dystrophin expression differ between muscles or even between different regions of the same muscle (Heier et al. 2023). This variability remains largely misunderstood but is thought to arise from variable regenerative capacities, levels of inflammation and quantity and quality of dystrophin (van de Bergen et al., 2014). Worldwide, the prevalence of DMD and BMD is of 1 in 3,500-5,000 boys (Walter & Reilich, 2017) and 1 in 18,500 boys (Bushby et al., 1991), respectively. While significant therapeutic advances have been made in the context of DMD, BMD is, generally speaking, significantly less studied and understood (Heier et al, 2023).

A major barrier in understanding disease mechanism has been the lack of appropriate BMD animal models. The first mouse model of BMD, referred as the *bmx* mouse model, was generated in 2022 by Heier and colleagues who used CRISPR-Cas9 technology to generate a 40,000 base pairs in-frame deletion of exons 45 to 47 (Heier et al., 2023). The *bmx* model seems to accurately recapitulate some of the phenotypes associated with BMD, including muscle weakness and heart dysfunction, and presents an intermediate phenotype between a wildtype and *mdx52* model, recapitulating DMD (Heier et al., 2023). Most of the phenotypes reported in their paper were measured in young *bmx* mice (10 to 15 weeks old), with the exception of cardiac phenotypes which were quantified in 18 months old mice.

In the context of BMD, mutations affecting different exons have been shown to correlate with a range of phenotypes and disease severities (Echigoya et al., 2018). Consequently, there is substantial value in modeling and characterizing mutations across various exons. This understanding led us to investigate another mutation situated in the second mutational hotspot— specifically, the deletion of exons 52 to 55. Here, we used CRISPR-Cas9 to generate a new *Dmd* del52-55 mouse model that harbors a typical BMD-like in-frame deletion. We demonstrate that, at an early age, *Dmd* del52-55 mice are mostly undistinguishable from wildtype mice. At 12 weeks of age, *Dmd* del52-55 mice show normal dystrophin expression and localization, muscle histology and mostly normal cardiac phenotypes. Functional tests also demonstrate that the truncated dystrophin expressed in *Dmd* del52-55 mice mostly maintains muscle function and protects against exercise-induced damage. However, some differences between *Dmd* del52-55 and wildtype mice seem to appear as mice grow older. Specifically, 52-week-old *Dmd del52-54* mice show reduced grip strength and contractile force after treadmill regimen. Therefore, our data suggest that a truncated protein is insufficient to protect against exercise-induced damage at 52 weeks. To further delineate the effects of the *Dmd* exons 52-55 deletion, we performed an unbiased transcriptomic analysis pre– and post-exercise and identified several differentially expressed pathways, including the heat shock and ubiquitin responses as well as the BMP signaling pathway, that could explain the fatigued phenotype observed at 52 weeks in our model. Overall, the characterization of this newly generated mouse model of BMD sheds some new light on BMD pathology and disease mechanisms.

## Results

### 1.1 Generation of a BMD in-frame *Dmd* del52-55 mouse model using CRISPR-Cas9

The deletion of exons 52 to 55 is a representative candidate mutation for generating a BMD-like in-frame deletion as it lies within one of the known mutational hotspots in the *DMD* gene (Figure 1A). To generate our new mouse model of a BMD-causing in-frame deletion, we designed four sgRNAs that remove a 213kb region spanning exons 52 to 55. Two sgRNAs targeting both introns 51 and 55 were designed to increase the frequency of editing events (Figure S1A). The sgRNAs and *Streptococcus pyogenes* Cas9 mRNA were delivered into fertilized zygotes through pronuclear injections and implanted in pseudopregnant females. The F0 mouse pups were genotyped for the presence of the predicted deletion junction, which was identified in 1 pup out of 25. Sanger sequencing also identified a deletion of an additional 5 base pairs upstream and downstream of the expected Cas9 cut sites (Figure S1B, C). The absence of exons 52 to 55 was confirmed through whole genome sequencing (Figure S1D). The splice sites were unaffected as validated by RT-PCR (S1E). These findings, thus, confirm the generation of an in-frame *Dmd* del52-55 mouse model.

**Figure 1:**
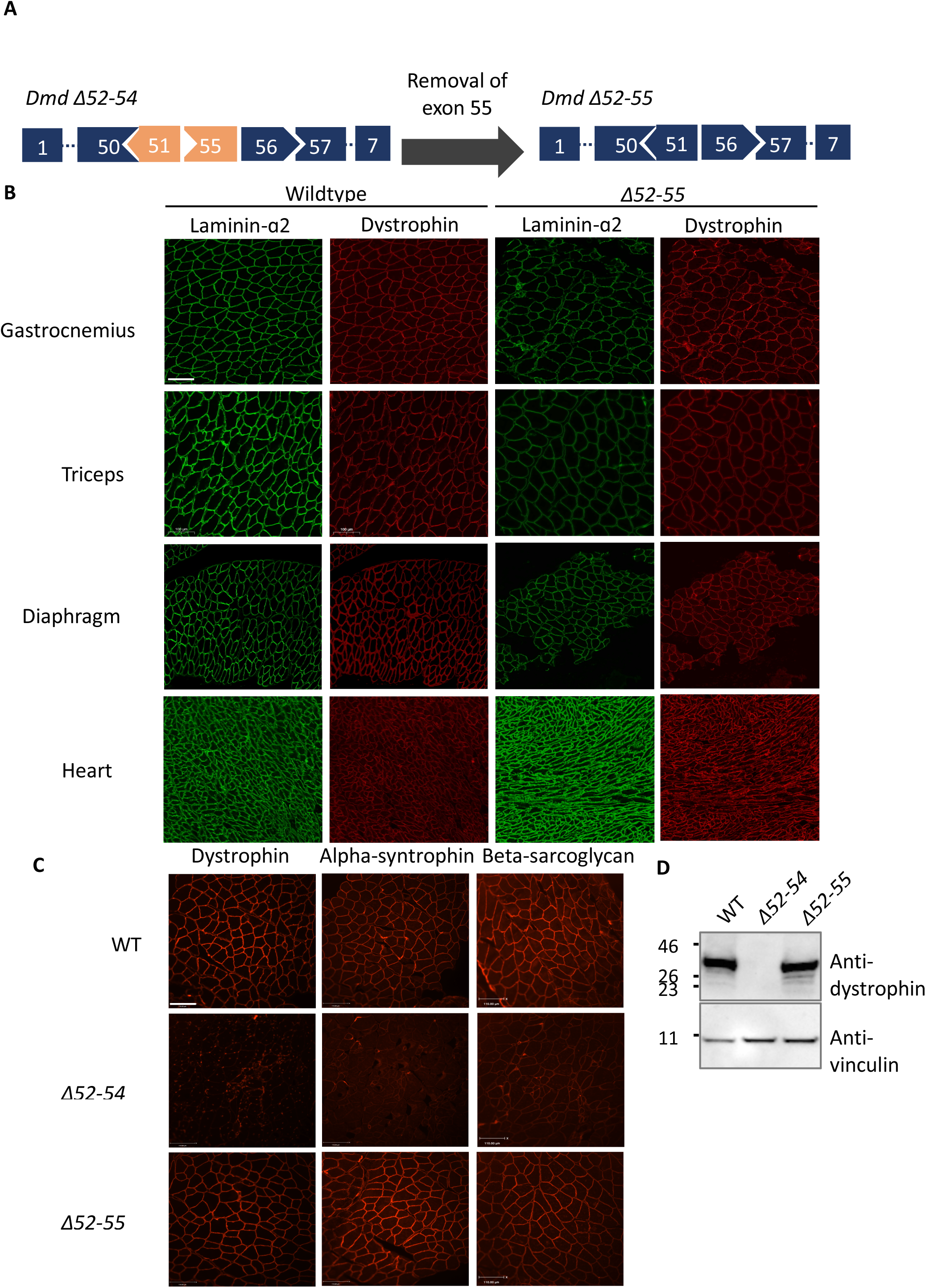
D*m*d del52-55 mice have WT-like levels of truncated dystrophin expression, which maintains normal DGC localization to the sarcolemma. (A) Schematic of the *DMD* gene showing the DMD-like deletion of exons 52 to 54 and the BMD-like in-frame deletion of exons 52 to 55. Both deletions occur in the central rod domain. (B) Immunofluorescence staining detected the sarcolemma using laminin-α2 (green) and the presence of dystrophin (red) in wild-type, *Dmd* del52-54 and *Dmd Δ52-54* cross sections of the gastrocnemius. Scale bar: 110 μm. (C) Immunofluorescence staining against dystrophin, alpha-syntrophin, nNOS and beta-sarcoglycan in WT, *Dmd* del52-55 mice and *Dmd* del52-54 mice. (D) Protein isolated from heart tissues showing dystrophin expression in the *Dmd* del52-55 mouse model but not in the DMD model *Dmd* del52*-54*. Vinculin was used as a loading control here.

### 1.2 *Dmd* del52-55 mice have WT-like dystrophin expression and localization at 12 weeks

Patients with DMD and BMD typically present with a lack or reduction of expression of dystrophin in muscle tissues (Duan et al., 2021; Bello et al., 2016; Heier et al., 2023). To understand the implications of a BMD-like in-frame deletion on the expression of dystrophin, we measured dystrophin expression levels in heart tissues in 12 weeks old mice. Of important note, our *Dmd* del52-55 mice were compared to *Dmd* del52-54 mice that harbor a complete DMD phenotype and WT mice. The results obtained for DMD and WT mice have already been published in an earlier volume of the same journal (see Wong et al. 2020). As shown in Figure 1B and 1D, expression levels were similar in *Dmd* del52-55 and WT mice in the tibialis anterior and gastrocnemius at 4 and 12 weeks old, respectively. Immunofluorescence staining indicates that dystrophin expressed in *Dmd* del52-55 localizes correctly to the sarcolemma in the gastrocnemius and recruits alpha-syntrophin, nNOS and beta-sarcoglycan, three components of the Dystrophin Glycoprotein Complex (Figure 1C). These findings indicate that 12 weeks old *Dmd* del52-55 mice have WT-like levels of truncated dystrophin expression, which maintains normal DGC localization to the sarcolemma at 12 weeks.

### 1.3 *Dmd* Del52-55 mice present WT-like muscle histology

Patients with DMD and BMD experience progressive muscle deterioration, which is characterized in Hematoxylin and Eosin-stained tissue sections by (1) immature myofibers containing centralized nuclei, (2) heterogeneity in muscle fiber sizes and (3) development of fibrotic tissue (Briguet et al., 2004). Here again, our *Dmd* del52-55 mice were compared to previously published results for *Dmd* del52-54 and WT mice (Wong et al., 2020) for easier comparison.

To determine the effects of the in-frame deletion of exons 52-55 on muscle degeneration and regeneration, we first quantified centralized nuclei in the gastrocnemius, triceps, and diaphragm of 12 weeks old mice. We observed less than 1% of myofibers with centralized nuclei in *Dmd* del52-55 mice, which is a level that is comparable to that observed in WT mice (Figure 2A, B). These results indicate that mice with a BMD-like in-frame deletion of exons 52 to 55 show normal levels of muscle turnover at 12 weeks of age.

**Figure 2:**
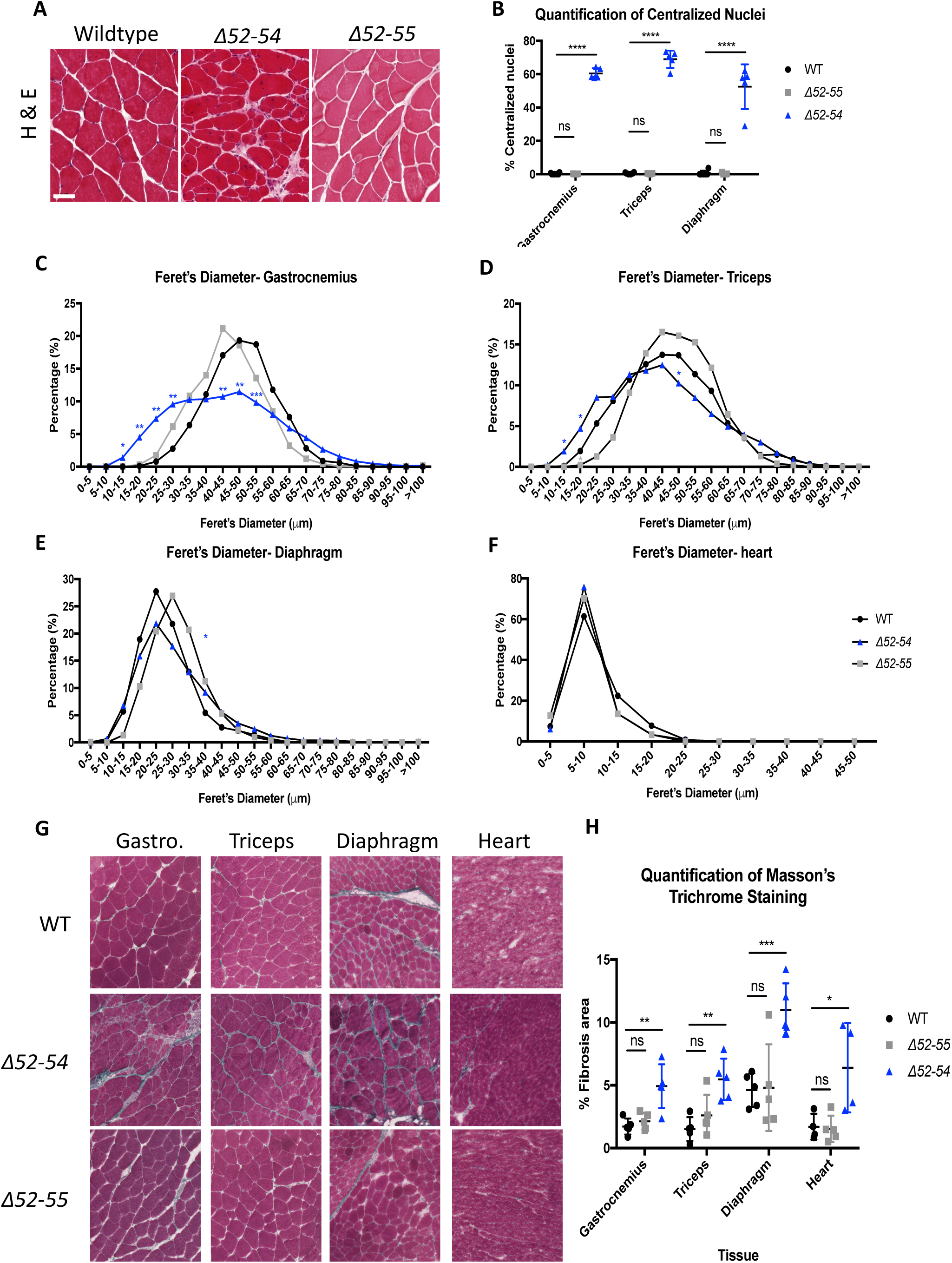
D*m*d del52-55 presents WT-like muscle histology. A) H&E staining of gastrocnemius cross sections from 12 week old wildtype, *Dmd Δ52-55*, *Dmd Δ52-54* and mice. Scale bar= 50μm. (B) Quantification of centralized nuclei in 12 weeks old mice WT (n=6) and *Dmd* del52-55 (n=5) mice. (B-C) Distribution of minimum Feret’s diameter quantified using H&E cross-sections from WT (n=4-5) and *Dmd* del52-55 (n=5) mice in the gastrocnemius (B) and triceps (C). (D) Masson’s trichome staining was performed on gastrocnemius, triceps, diaphragm and heart tissues of 12 weeks old wildtype (n=4-5) and *Dmd* del52-55 (n=5) mice and the fibrotic area was quantified. Scale bar: 250 μm. All data is represented as the mean ± S.D. Statistical analyses were performed with Student’s t-test. P* < 0.05, P** < 0.01, P*** < 0.001, P**** < 0.0001.

The size of cross sections of myofibers was then quantified based on minimal Feret’s diameter in the gastrocnemius, triceps, diaphragm and heart of 12 weeks old mice. Results indicate that myofibers were comparably homogeneous in size in WT and *Dmd* del52-55 mice across tissues, which suggests that normal muscle degeneration and regeneration is occurring (Figure 2C, D, E, F).

Levels of fibrotic tissue were quantified using Masson’s trichrome staining on the gastrocnemius, triceps, diaphragm and heart of 12 weeks old mice. Results suggest that fibrotic changes are not observed in *Dmd* del52-55 mice (Figure 2G, H).

Taken together, these results indicate that a BMD-like in-frame deletion of exons 52 to 55 does not affect muscle histology and levels of muscle degeneration and regeneration. At 12 weeks of age, we, therefore, see WT-like muscle histology in mice expressing a truncated dystrophin protein.

### 1.4 *Dmd* del52-55 mice have WT-like cardiac phenotypes

Patients with Duchenne Muscular Dystrophy and Becker Muscular Dystrophy tend to present with progressive cardiac hypertrophy, arrhythmia or dilation of the ventricles due to the reduction of dystrophin expression in cardiac tissues (Hermans et al., 2010; Heier et al., 2023).

Echocardiography results show that our *Dmd* del52-55 model presents with similar posterior wall thickness during systole, systolic volume, diastolic volume and end systolic diameter as WT mice throughout 12 to 52 weeks of age (Figure 3A, C, D, E). We see a small, yet significant increase in posterior wall thickness during diastole in *Dmd* del52-55 compared to WT mice at 12 weeks of age (Figure 3B) and a decreased end diastolic diameter at 28 weeks of age (Figure 3F). In Figures 3A, C and D, results from our *Dmd* del52-55 mice were compared to previously published results for *Dmd* del52-54 and WT mice (Wong et al., 2020) for easier comparison.

**Figure 3:**
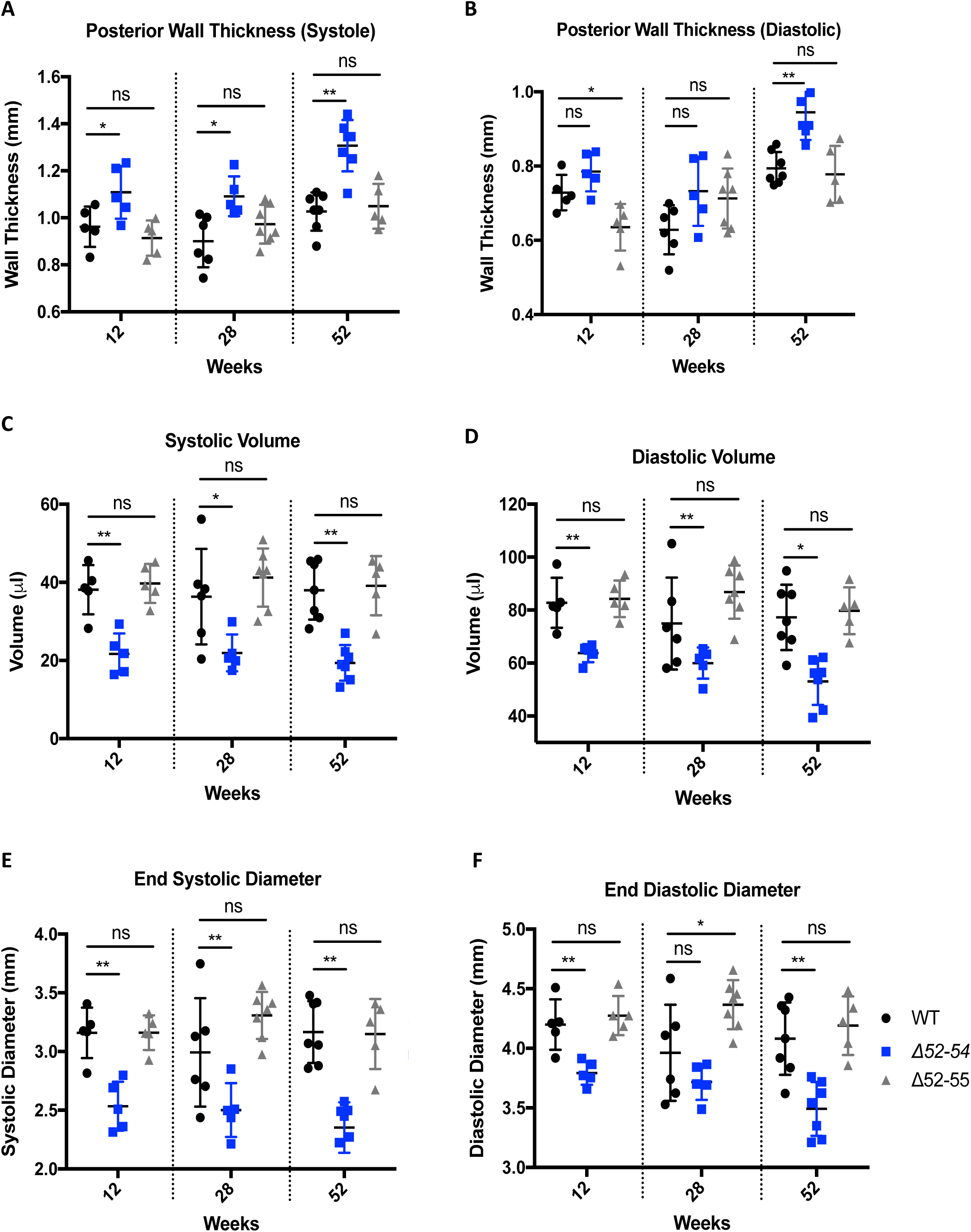
D*m*d del52-55 mice maintain mostly wildtype cardiac anatomical features throughout 12 to 52 weeks of age. The heart in WT and *Dmd* del52-55 mice were analyzed using echocardiography to determine (A) left ventricular posterior wall thickness during systole and (B) diastole, (C) left ventricular systolic and (D) diastolic, as well as (E) left ventricular end systolic and (F) diastolic diameter. All data is represented as ± S.D. Statistical analyses were performed with Student’s t-test. P* < 0.05, P** < 0.01.

*Nppa* expression levels were measured in *Dmd* del52-55 mice as a proxy for cardiac stress. Results indicate that *Dmd* del52-55 mice have similar levels of *Nppa* expression to WT mice (Figure S2A). Heart rate in *Dmd* del52-55 mice was also similar to WT throughout 12 to 52 weeks of age, indicating an absence of tachycardia in mice expressing the truncated dystrophin protein (Figure S2B). The ejection fraction and fractional shortening were similar between *Dmd* del52-55 and WT mice (Figure S2C and D). Overall, *Dmd* del52-55 mice expressing a truncated dystrophin protein maintain mostly normal cardiac parameters similar to wildtype mice, indicating limited cardiac stress in these mice.

### 1.5 Truncated dystrophin expressed in *Dmd* del52-55 mice maintains muscle function and protects against exercise-induced damage at 12 weeks of age

In Duchenne and Becker Muscular Dystrophies, the presence of fibrosis typically impairs muscle function and regeneration capacity (Capitanio et al., 2020). To assess the ability of the truncated dystrophin protein expressed in *Dmd* del52-55 mice to protect against exercise-induced muscle damage, several functional tests were performed, including grip strength and tetanic force after treadmill regimen. At 12 weeks of age, *Dmd* del52-55 mice showed wildtype-like grip strength (Figure 4A). *Dmd* del52-55 mice showed no areas with damaged fibers before or after the treadmill regimen (Figures 4B and C). Tetanic force was also measured in sedentary and exercised *Dmd* del52-55 mice, which indicates a similar response to exercise between *Dmd* del52-55 and WT mice, namely an increase in muscle strength after exercise (Figure 4D). Although *Dmd* del*52-55* expresses a shorter dystrophin protein, there is no detriment to grip strength function at 12 weeks of age. Similarly to full-length dystrophin, the shorter dystrophin protein protects from muscle stimulation while allowing enhanced muscle strength after exercise.

**Figure 4:**
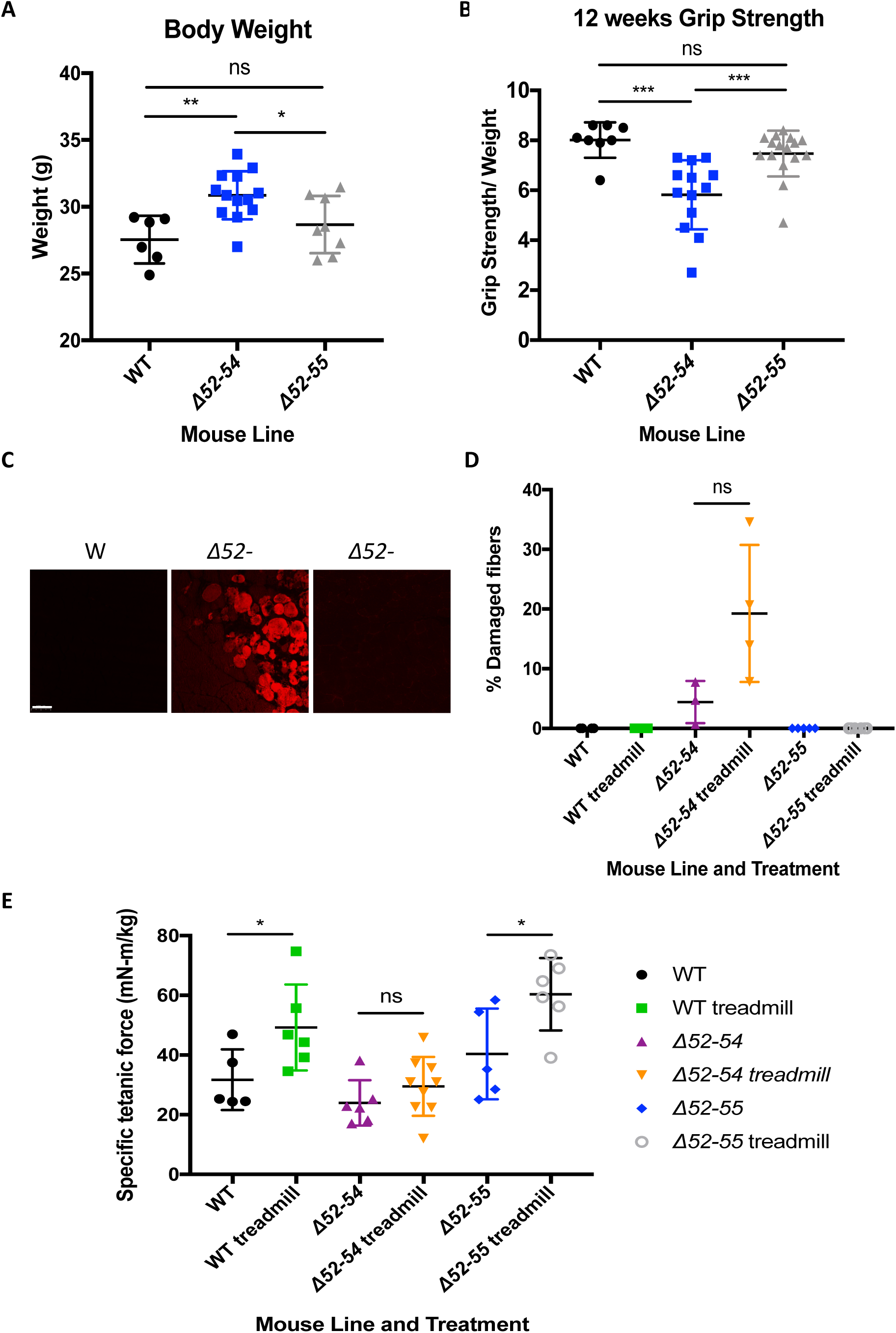
Truncated dystrophin expressed in *Dmd* del52-55 is not detrimental to muscle function and protects from muscle aggravation at 12 weeks. (A) Forelimb and hindlimb grip strength of wildtype (n=8) and *Dmd* del52-55 (n=16) mice. (B) Gastrocnemius cross-sections from exercised wildtype and *Dmd* del52-55 mice injected with Evan’s blue dye (EBD). (C) EBD positive myofibers in gastrocnemius of sedentary and exercised wildtype and *Dmd* del52-55 mice were quantified. Scale bar = 100 μm. (D) Tetanic force was measured in sedentary and exercised mice from wildtype and *Dmd* del52-55 mice. All data is represented as ± S.D. Statistical analyses were performed with Student’s t-test. P* < 0.05, P** < 0.01, P*** < 0.001.

### 1.6 Truncated dystrophin expressed in *Dmd* del52-55 mice is not sufficient to maintain normal muscle function at 52 weeks of age

Although the truncated dystrophin protein expressed in *Dmd* del52-55 seems to mostly maintain muscle function at 12 weeks of age, a few differences between *Dmd* del52-55 and wildtype mice (including smaller fibers observed in the gastrocnemius in *Dmd* del52-55 mice and some cardiac phenotypes that are different between the two groups), prompted us to investigate muscle function in older, 52-week-old mice.

At 52 weeks of age, forelimb grip strength was lower in *Dmd* del52-55 mice compared to wildtype mice after treadmill exercise (Figure 5A). *Dmd* del52-55 mice also showed a decrease in contractile force at 75, 100 and 125 Hz after exercise compared to wildtype mice (Figure 5C). While differences were observed in grip strength and contractile force at 52 weeks of age, performances in open field parameters were similar between *Dmd* del52-55 and wildtype mice pre– and post-exercise (Figure 5D). This data suggests that a truncated dystrophin protein may not be sufficient to maintain fully wildtype muscle function with age.

**Figure 5:**
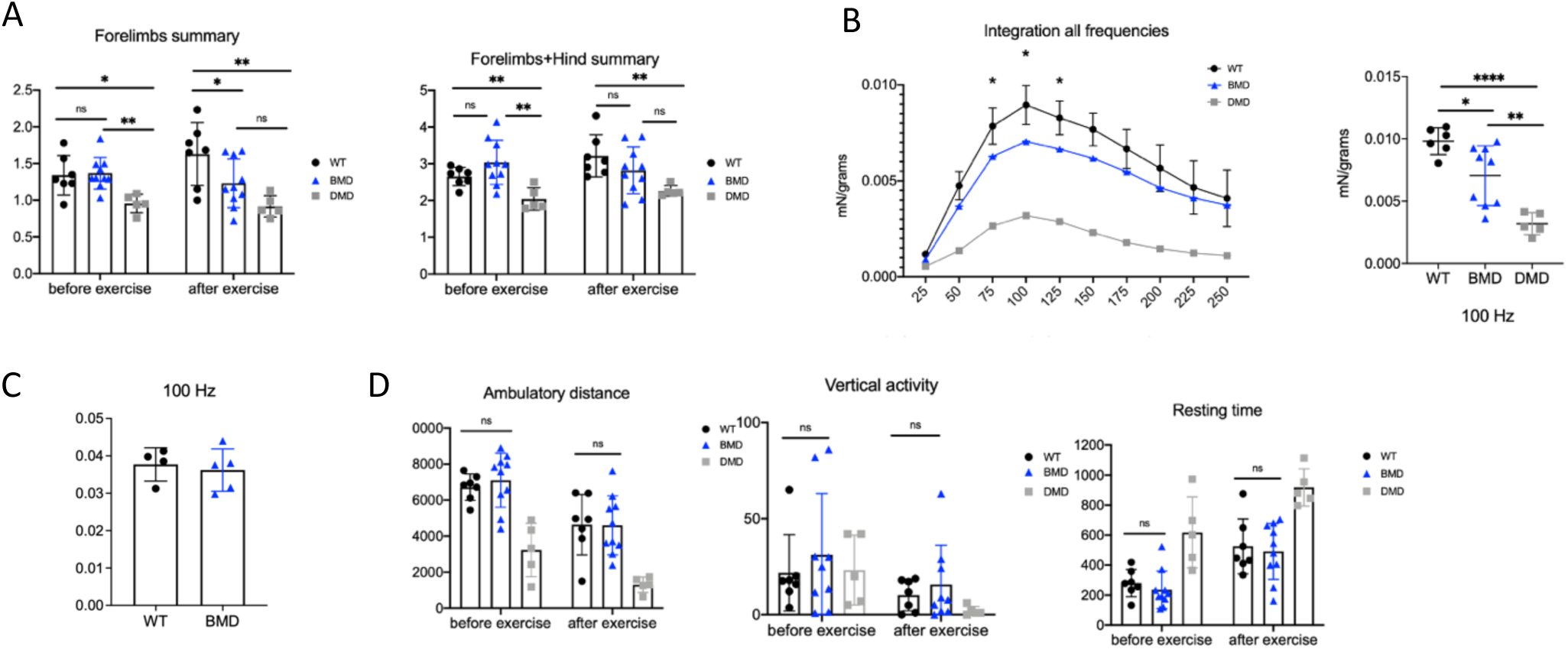
D*m*d del52-55 mice show impaired muscle function at 52 weeks of age. (A) Forelimb and combined forelimb and hindlimb grip strength in wildtype (n=7) and *Dmd* del52-55 (n=8) mice. (B) Contractile force in WT and *Dmd* del52-55 mice measured after treadmill exercise. (C) Contractile force in sedentary WT, *Dmd* del52-54 and *Dmd* del52-55 mice. (D) Open field performances in ambulatory distance, vertical activity and resting time in WT (n=7) and *Dmd* del52-55 (n=9-10) mice. All data is represented as ± S.D. Statistical analyses were performed with Student’s t-test. P* < 0.05, P** < 0.01, P*** < 0.001, P**** < 0.0001.

### 1.7 Exercise induces differential gene expression in BMD mice at 52 weeks

Given the lower grip strength and contractile force observed in 52-week-old-exercised *Dmd* del52-55 mice, we hypothesized that exercise aggravated the fatigue phenotype by inducing transcriptional changes involved in muscle regeneration and degeneration. We performed RNA-Seq on the gastrocnemius of 52-week-old sedentary and exercised BMD-like mice (*Dmd* del52-55), DMD mice (*Dmd* del52-54) and WT mice. *Dmd* del52-54 mice were used as a reference and behaved as expected, with a large number of differentially expressed genes compared to WT mice both pre– and post-exercise (Figure S3). While only four genes were differentially expressed between the BMD-like and WT cohorts of sedentary mice, 53 genes were up-or down-regulated in the exercised BMD-like mice compared to WT. Some of these genes, including *Hspa1a* and *Hspa1b*, two protein folding chaperones, were substantially up-regulated (97– and 50-fold change, respectively). These data suggest that exercise induce significant transcriptional changes in in *Dmd* del52-55 mice (Figure 6A).

**Figure 6:**
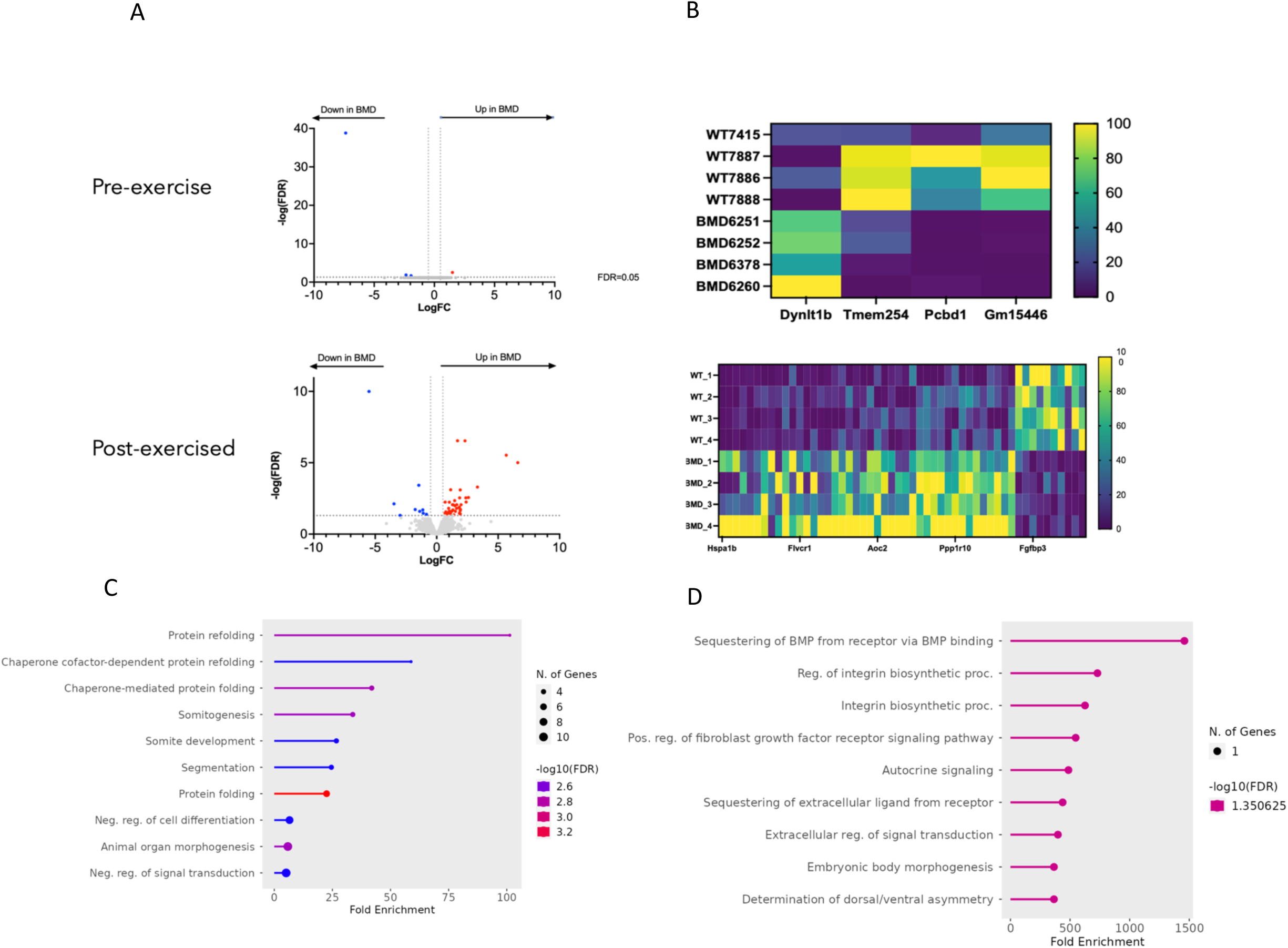
Exercise induces differential gene expression in the gastrocnemius of 52-week-exercised *Dmd* del52-55 mice. (A) Volcano plots showing genes that are upregulated (in red) and downregulated (in blue) in *Dmd* del52-55 mice (n=4) compared to WT mice (n=4) before and after exercise. (B) Heat maps showing genes that are up– and down-regulated in *Dmd* del52-55 mice (n=4) compared to WT mice (n=4) before and after exercise. (C, D) Gene Ontology Analysis showing the distribution of GO terms significantly up-(C) and down-regulated (D) between *Dmd* del52-54 and wildtype.

To better understand which pathways were involved, we performed Gene Ontology and Gene Set Enrichment analyses (see Figure S3) for each cohort. As expected for DMD mice, upregulated gene pathways included those involved in inflammation, wound healing and calcium signaling (dysfunction of calcium-dependent proteases), while downregulated pathways included cytoskeletal organization (especially in exercised mice), the release of sequestered Ca2+ ions and calcium binding (acting upstream of the actin cytoskeleton reorganization), muscle and heart contraction and development (Figure 6B). We then identified differentially expressed in the BMD exercised mice (*Dmd* del52-55) compared with WT exercised mice (Figure 6C). Upregulated pathways included protein folding, clearance of protein aggregation, heat shock and ubiquitin responses and the organization of the cytoskeleton, while downregulated pathways included BMP signaling or belong to the Gm family of long non-coding RNAs (Figure 6C). Taken together, these results suggest that gene expression is influenced by exercise in BMD mice leading to the activation of catabolic pathways in skeletal muscles.

## Discussion

Here, we generate and characterize a new mouse model of BMD, in which CRISPR-Cas9 technology was used to generate a BMD-like in-frame deletion of exons 52 to 55. This deletion is located in one of the two mutational hotspots in the *DMD* gene and is, therefore, likely to be relevant to patients with BMD. To our knowledge, this is the first mouse model of BMD in which phenotypes other than those affecting the heart were studied over a period of 52 weeks. We demonstrate that, at an early age, *Dmd* del52-55 mice are mostly identical to wildtype mice: at 12 weeks of age, *Dmd* del52-55 mice show normal dystrophin expression and localization, normal muscle histology and mostly normal cardiac phenotypes. Functional tests also demonstrate that the truncated dystrophin protein expressed in *Dmd* del52-55 mice mostly maintains muscle function and protects against exercise-induced damage. This is consistent with the fact that exons 52 to 55 are within the central rod domain, thus leaving the essential actin binding domain, cysteine-rich domain and C terminal domain unaffected (Sun et al., 2020).

However, some differences between *Dmd* del52-55 and wildtype mice seem to appear as mice age and are aggravated by exercise. Exercised 52-week-old *Dmd* del52-55 mice show reduced forelimb grip strength and contractile force compared to WT mice. Our unbiased transcriptomic data suggests that this fatigued phenotype is associated with a change in gene expression that appears to be induced by exercise. Looking more closely at the genes that are differentially expressed in *Dmd* del52-55 exercised mice may provide some preliminary insights into the mechanisms underlying this fatigued phenotype. Gene Ontology and Gene Set Enrichment analyses revealed that genes involved in heat shock and ubiquitin responses are upregulated in our newly generated BMD model. This trend has also been observed in DMD mice (Nishimura & Sharp, 2005; Bornman et al. 1995; Wattin et al. 2018). De Paepe and colleagues (2012) also note that heat shock proteins and chaperones balance muscle regeneration and destruction in DMD patients. They analyzed muscle biopsies from seven DMD patients and conclude that heat shock proteins are upregulated in regenerating, atrophic muscles as well as in macrophages and cytotoxic T-cells invading non-necrotic muscle fibers (De Paepe et al. 2012). A similar mechanism could exist in BMD, where the upregulation of heat shock proteins and chaperones would be present in both regenerating fibers and immune cells. Further work is required to determine whether this hypothesis is valid or whether the heat shock and ubiquitin responses are only observed in a single cell type, thereby promoting either protection or destruction.

Gene Ontology and Gene Set Enrichment analyses also identified genes that are downregulated in our newly generated BMD model, namely genes involved in BMP signaling and pertaining to the Gm family of long non-coding RNAs. In the context of BMP signaling, our RNA-Seq data reveals that the expression of *Grem2*, a BMP antagonist, is downregulated in our BMD-like *Dmd* 52-55 mice. Homeostatic BMP signaling is integral to muscle differentiation and regeneration capacity. Downregulation of *Grem2* has been observed in DMD patients (Pescatori et al. 2007): Pescatori and colleagues (2007) analyzed genome-wide gene expression profiles of 19 DMD patients and identified *Grem2* as one of the top 18 differentially expressed genes in muscles. Shi and colleagues also observed inefficient differentiation of myoblast in DMD mice models (Shi et al. 2011; Shi et al. 2013). Here, it could be hypothesized that the downregulation of *Grem2* leads to the upregulation of BMP signaling, which, in turns, prevents muscle cell differentiation and muscle regeneration in our BMD-like mice. Further work is required to validate this hypothesis and determine whether this mechanism significantly contributes to BMD pathology. Furthermore, although it is known that some of the genes that belong to the Gm family of long non-coding RNAs mentioned earlier are involved in RNA Pol II DNA binding activity, little is known about this gene family overall. As such, further work would be necessary to shed light on their contribution to BMD pathology.

As noted by Heier and colleagues (2023), it is important to generate more models of Becker Muscular Dystrophy in view of comparing them and identifying which dystrophin isoforms (and internal deletions) are associated with what phenotype. The latter is important for at least three reasons: (1) to provide better predictions about disease severity and phenotype to patients and their families, (2) to identify the effects of treatments for different BMD mutations, (3) to determine which DMD patients are more amenable to exon skipping and predict the extent to which the dystrophic phenotype would be altered using this approach (Bello et al., 2016). While deletions in the N-or C-terminal domains typically result in a DMD phenotype, deletions in the rod domain of dystrophin may have variable effects based on the structural “phase” between spectrin repeats (Bello et al., 2016; Duchêne et al., 2018). It has been observed that in-frame mutations in exons 50-51 typically result in a milder BMD phenotype while mutations in exons 45-53 (excluding exons 50-51) usually lead to a more severe BMD phenotype (Bello et al., 2016). Little is known about mutations in the remaining part of the mutational hotspot, namely exon 53 to 55. Our newly generated model of BMD addresses this knowledge gap. Results obtained with our *Dmd* del52-55 mouse model can be compared with those observed in the *bmx* mouse model (deletion of exon 45 to 47) generated by Heier and colleagues (2023), as shown in Figure 7. Overall, it appears that the BMD phenotype is more severe in the *bmx* model than in our *Dmd* del52-55 model. This is consistent with our knowledge of dystrophin: the deletion of exons 45 to 47 is predicted to disrupt spectrin repeats and nNOS binding (Heier et al., 2023), thereby significantly disrupting the DGC complex. In the future, we hope that such comparisons can be made between different models of BMD in order to gain insights on the functionality of different dystrophin isoforms.

**Figure 7:**
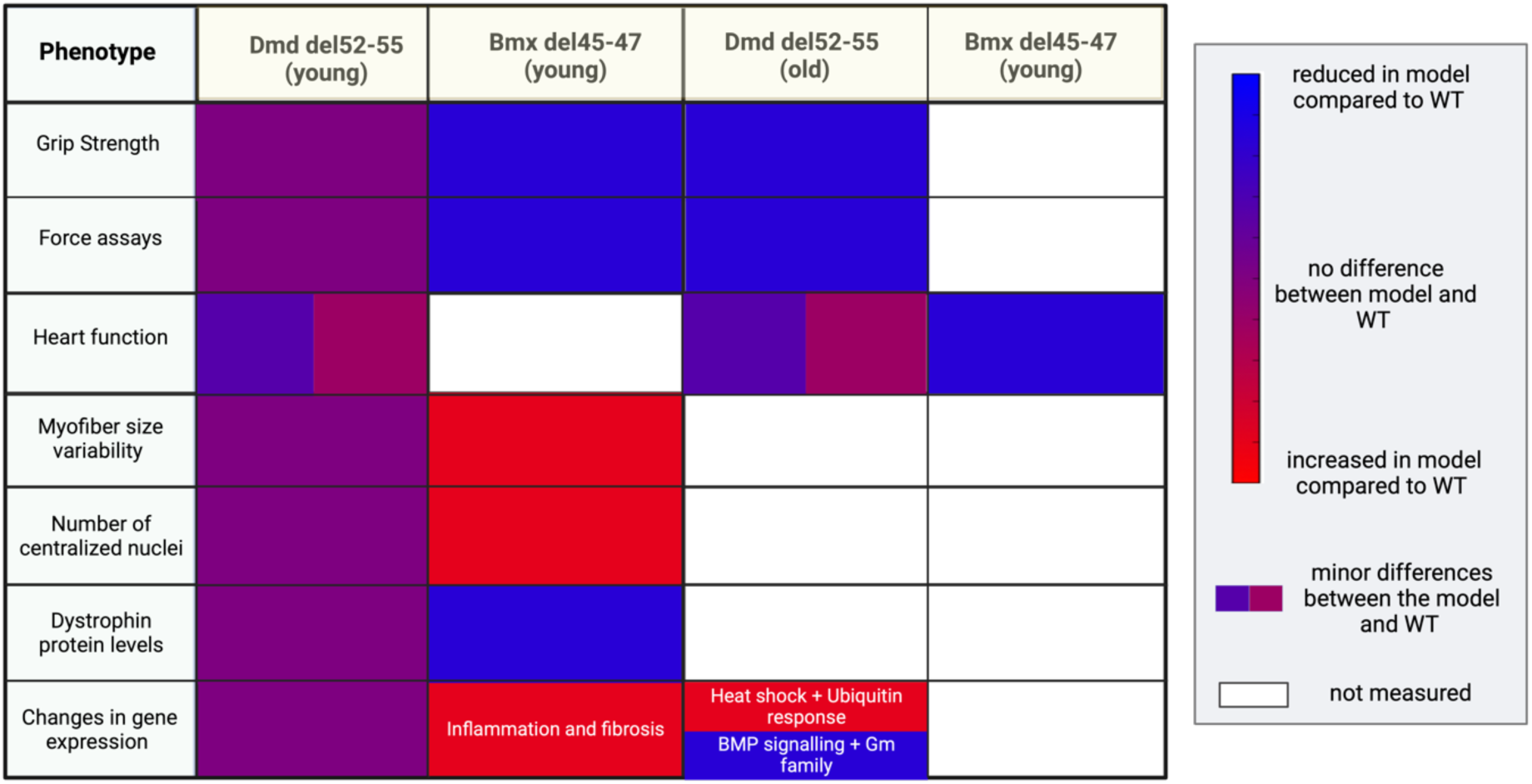
Comparisons of phenotypes observed in the *bmx* model and our *Dmd* del52-55 model.

Treatment options for patients with BMD are very limited, even more so than for patients with DMD. This issue is exacerbated by the fact that very few clinical trials are currently being conducted for patients with BMD. Glucocorticoid treatment, which is the most widely used and accessible treatment option for DMD patients, is rarely used in BMD patients since the risk-benefit profile of this treatment for a longer BMD disease course remains unclear (Heier et al., 2023). Glucocorticoids have anti-inflammatory and immunosuppressive properties and were shown to delay loss of ambulation in patients with muscular dystrophies (Bello et al. 2015). However, glucocorticoids come with significant side effects, such as weight gain, adrenal suppression and mood imbalances (Heier et al., 2023). Recently, Vamorolone, a new dissociative anti-inflammatory with improved safety profile has been approved by the FDA (Heier et al., 2019). We hope that BMD mouse models and natural history studies will continue to be used in conjunction to further advance treatments and identify which therapeutic option is best according to the underlying mutation exhibited in individual patients.

## Materials and Methods

### 1. sgRNA design

sgRNAs specific for the *Strepcoccus pyogenes* Cas9 system were designed using benchling.com (Table 1). All sgRNAs were synthesized following Gertsenstein and Nutter (2018).

**Table 1.**
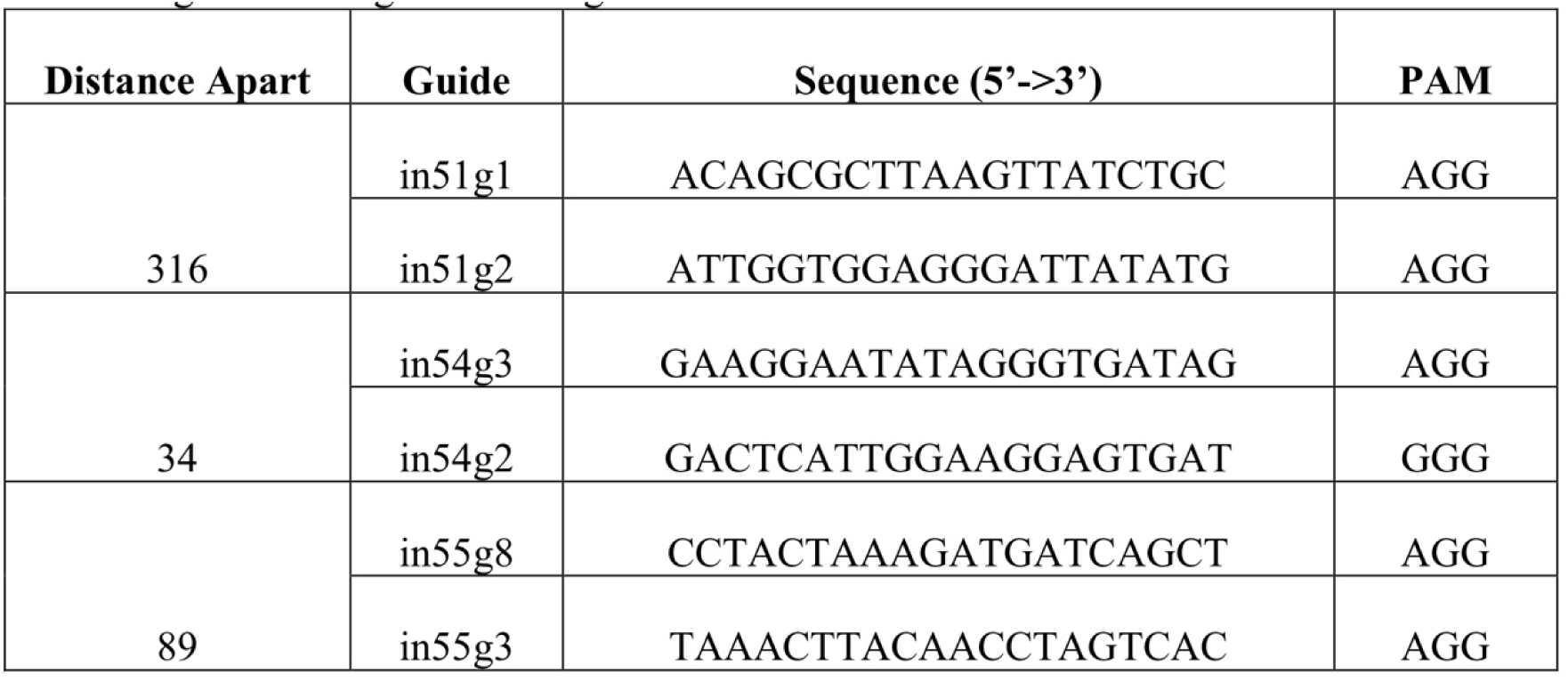
sgRNAs designed for the generation of *Dmd Δ52-54* and *Dmd Δ52-55 mouse* lines.

### 2. Pronuclear injections

3-4 weeks old C57BL/6J (Jackson) females were used as embryo donors. Pseudopregnant surrogates were CD-1 (ICR) (Charles River) females. All procedures were conducted following Behringer et al. 2014. Briefly, embryo donors were superovulated and mated overnight with males. Successfully mated females were selected and the oviducts were dissected to isolate fertilized zygotes. Zygotes were then subjected to pronuclear microinjections with sgRNAs and Cas9 mRNA. Each injection was comprised of 10ng/uL of each sgRNA and 20ng/uL of Cas9 mRNA. Injected embryos were transferred into the oviducts of the surrogate females. Mice were backcrossed for three generations and then intercrossed before used for analysis. Only males were used in this study.

### 3. DNA extraction and analyses

Mouse tails were biopsied at 2 weeks old, and DNA was extracted using a DNAeasy Blood and Tissue kit (Qiagen). One microliter of the extracted DNA was used for PCR with DreamTaq polymerase (Thermo Fisher Scientific) and the primers shown below in Table 2.

**Table 2:**
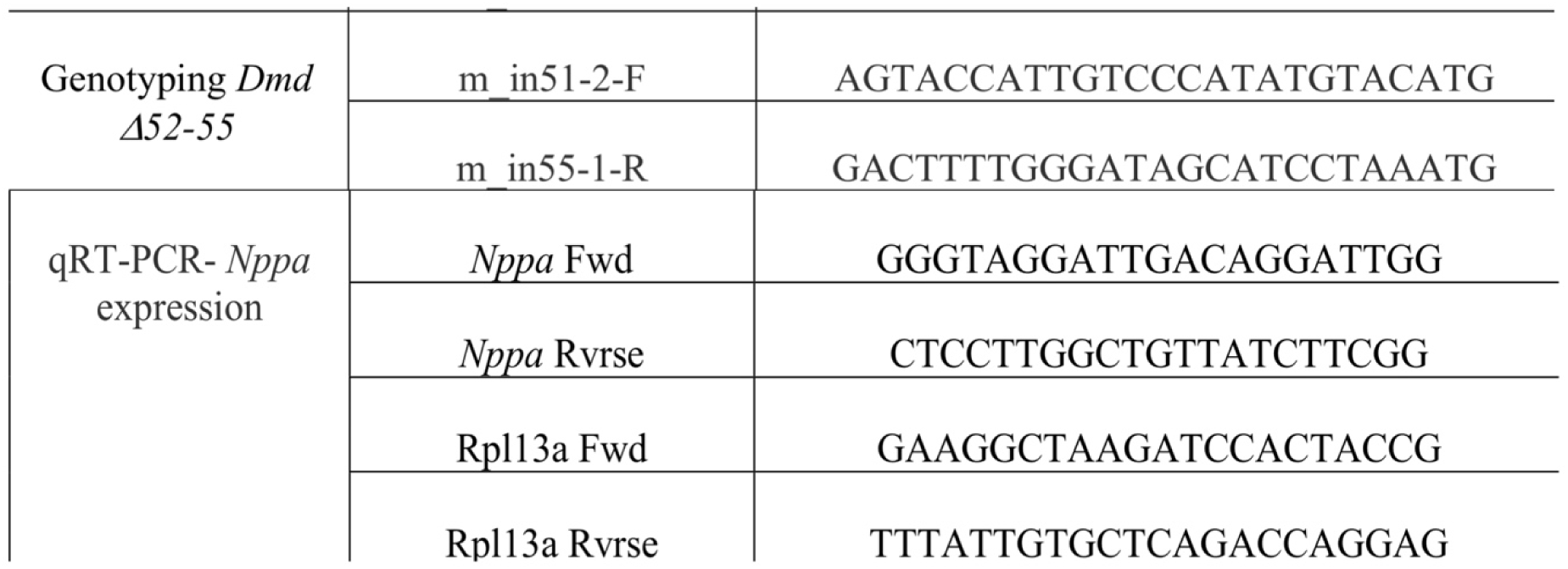
Oligonucleotides utilized for molecular analyses.

### 4. Whole genome sequencing

DNA extracted from mouse tails was used for whole genome sequencing (WGS), which was performed using the Illumina HiSeq X system (San Diego, CA, USA) by The Centre for Applied Genomics (TCAG) at the Hospital for Sick Children. A Qubit Fluorometer High Sensitivity Assay (Thermo Fisher Scientific, Waltham, MA, USA) was used to measure DNA yield and the Nanodrop (Thermo Fisher Scientific) OD260/OD280 ratio was used to confirm DNA purity. 400 ng of DNA sample was used for library preparation using the Illumina TruSeq PCR-free DNA Library Prep Kit. DNA was then sonicated into 350-bp fragments. A-tailed and indexed TruSeq Illumina adapters were ligated to end-repaired sheared DNA fragments before library amplification. Libraries were analyzed using Bioanalyzer DNA High Sensitivity chips (Agilent Technologies, Santa Clara, CA, USA) and quantified using qPCR. The libraries were loaded in equimolar quantities and pair-end sequenced on the Illumina HiSeqX platform to generate 150-bp reads. Integrative Genomics Viewer (IGV) version 2.8.2 was used for analysis with GRCm38/mm10 as the murine reference genome.

### 5. RNA extraction and sequencing

Mouse gastrocnemii were sectioned into 30-um slices, collected in 1.4 mm Zirconium Beads tubes (OPS Diagnostic) and homogenized using a MNagNA Lyser (Roche Diagnostic). RNA extraction was performed using Trizol reagent (Thermo Fisher Scientific) as per the manufacturer’s protocol. Following RNA extraction, RNA was quantified using a Qubit RNA HS assay (Thermo Fisher Scientific). Sequencing was performed by TCAG (Hospital for Sick Children, Toronto, Ontario) using the Illumina HiSeq 2500 system, which produces 150-bp paired-end reads. Raw transcript reads were aligned to the GRCm38/mm10 mouse reference genome using STAR aligner, v.2.6.0c. (https://github.com/alexdobin/STAR). HTSeq v.0.6.1p2 was used to determine the absolute number of read counts for each gene. Normalization and differential expression analysis were completed using the R package DESeq2 v.1.26.0s package (https://bioconductor.org/packages/release/bioc/html/DESeq2.html). Initial minimal filtering of 10 reads per gene for all samples was applied to the datasets. Differentially expressed genes were defined as having a FDR < 0.05 and FC > 1.5 (for upregulated genes) or < 0.6 (for downregulated genes. Gene ontology analysis of the DEGs was performed using ShinyGo and Gene Set Enrichment Analysis was performed using the UC San Diego GSEA software (https://www.gsea-msigdb.org/gsea/index.jsp). Heat maps and volcano plots were made using GraphPad Prism v9.

### 6. Animal husbandry

All mice were housed at The Centre for Phenogenomics (TCP) in Toronto, Canada under the environmental regulations of 12 hour light/dark cycle with food and water provided in individual units (Techniplast). All procedures were conducted in accordance with the Animals for Research Act of Ontario and the Guidelines of the Canadian Council on Animal Care. Animal protocols were performed at The Center for Phenogenomics (TCP) Toronto and were reviewed and approved by the local Animal Care Committee.

### 7. Functional tests

Forelimb and hindlimb grip strength tests were conducted by The Centre for Phenogenomics in Toronto based on the TREAT-NMD SOP DMD_M.2.2.001 protocol. 12-, 28– and 52-week-old male C57BL/6J and Dmd Δ52-54 were placed over the grid of the grip strength meter (Bioseb). Forepaws and hindpaws were allowed to attach to the grid before pulling the mouse back by the tail, and the maximal grip strength value of the mouse was measured. The grip strength test was done in triplicates, where the average grip strength value was normalized by the mouse body weight.

For the echocardiography, male mice were scanned using the Vevo2100 ultrasound machine (VisualSonics, Toronto, Canada) with a 30 MHz transducer as described previously (Zhou et al., 2005). All mice were scanned under 1.5% isoflurane anesthesia for 20-30 min with careful monitoring of the body temperature to maintain it at 37-38°C (TREAT-NMD DMD_M.2.2.003). All measurements were conducted using the cardiac package of the Vevo 2100 v1.6.0 software.

For the open field test, mice were placed in the frontal center of a transparent Plexiglas open field (41.25 cm × 41.25 cm × 31.25 cm) illuminated by 200 lux. The VersaMax Animal Activity Monitoring System recorded activity in the center and periphery of the open field arena for 20 min per animal.

In vivo tibialis anterior contraction tests were performed as previously described (Kemaladewi et al., 2019). Briefly, contractile activity was measured using the 1300A: 3– in-1 Whole Animal System and analyzed using the Dynamic Muscle Analysis 5.5 and 5.3 high-throughput software (Aurora Scientific). The mice were anaesthetized using ketamine-xylazine solution at 100 mg/kg and 10 μl mg/kg to body weight, respectively, through intraperitoneal injection. Percutaneous electrodes were placed in the tibialis anterior and contractile output was measured.

For treadmilling exercises, 6-week-old C57BL/6J, Dmd del52-55 and Dmd del52-54 mice were placed on a treadmill (Columbus) which was set up at a downhill angle of 15° as previously described (Anderson, 2006). The mice were run for 10 minutes at 12 m/min for 3 consecutive days. All exercised and sedentary mice were injected with EBD after the second day of exercise and in vivo contractile tests were performed after treadmill exercise on all mice after the third day of exercise. Muscle tissues were dissected 24 hours post EBD injection.

### 8. Preparation and injection of Evan’s blue dye and assessment

Evan’s Blue Dye (EBD) (Fisher Scientific) stock was prepared at 1% (w/v) in 1× phosphate-buffered saline (PBS– and filtered through Millex®-GP 0.22 μm filter (Millipore, Bedford, MA, USA) (Hamer, 2002). Muscles dissected from mice injected with EBD were sectioned at 8 μm and mounted with ProLong Gold Antifade Mountant (Thermo Fisher Scientific) EBD saturated myofibers fluoresce at 590nm, and were scanned with the 3DH Panoramic Slide Scanner at the Imaging Facility at the Hospital for Sick Children and images were acquired with CaseViewer (3DHISTECH). Quantification was performed using ImageJ 1.52a software and percentages of damaged fibers were calculated as follows: % damaged fibers = EBD area / total tissue area.

### 9. Tissue processing

Shortly following cervical dislocation, mouse hearts were arrested in diastole through direct KCl (1 M KCl in PBS) injection. All muscles dissected were frozen in cooled isopentane in liquid nitrogen as previously described (Nelson et al., 2016).

### 10. Histological staining (H&E)

All muscles were sectioned at 8 μm for histological staining. Hematoxylin and Eosin (H&E) staining was conducted using a standard protocol (Fischer et al., 2008). H&E slides were then scanned using the 3DH Pannoramic Slide Scanner by the Imaging Facility at the Hospital for Sick Children. CaseViewer (3DHISTECH) was used for image acquisition. Centralized nuclei were quantified using ImageJ 1.52a software from a total of 300 myofibers.

### 11. Masson’s trichome staining

Trichrome staining was performed at the Pathology lab at The Centre for Phenogenomics, Toronto (TREAT-NMD SOP MDC1A_M.1.2.003). Trichrome-stained slides were scanned using a Hamamatsu Nanozoomer and analyzed using the NDP.view2 Viewing Software. Three frames containing at least 300 fibers each were used for fibrosis quantification using ImageJ 1.51 software.

### 12. Immunofluorescence staining and analysis

8 μm sections of muscle tissues were used for immunofluorescence staining. Sections were fixed in ice-cold methanol and blocked with blocking buffer (3% normal goat serum and 0.2% BSA in PBS). Primary antibodies were incubated overnight at 4°C. We used the following primary antibodies: rabbit polyclonal anti-dystrophin (abcam15277; Abcam; 1:200), rabbit polyclonal anti-syntrophin-α 1 (ab11187; Abcam; 1:600), rabbit polyclonal anti-β-sarcoglycan (ab222241; Abcam; 1:100) and rat monoclonal anti-laminin-2 (α2 chain; 4H8-2; Sigma Aldrich; 1:500). Secondary antibodies used were goat polyclonal anti-rabbit Alexa Fluor 594-conjugated (Thermo Fisher Scientific; 1:250) and goat polyclonal anti-rat Alexa Fluor 488-conjugated (Thermo Fisher Scientific; 1:250) antibodies. Tissues were incubated at room temperature with secondary antibodies for 2 hours. For sections stained with anti-laminin-α2, slides were scanned as described previously (Kemaladewi et al., 2017). Feret’s diameter was quantified (TREAT-NMD SOP DMD-M1.2.001) using Open-CSAM in the ImageJ 1.51 software (Desgeorges et al., 2019).

### 13. Western Blotting

We extracted proteins from homogenized mouse tissue by using a 1:1 solution of RIPA homogenizing buffer (50 mM Tris-HCl pH 7.4, 150 mM NaCl and 1mM EDTA) and RIPA double-detergent buffer (2% deoxycholate, 2% NP-40, 2% Triton X-100 in RIPA homogenizing buffer) supplemented with protease inhibitor cocktail (Roche), as described previously (Kemaladewi et al., 2019). The concentration of each protein was quantified using a Pierce BCA protein assay kit (Thermo Fisher Scientific). 15μg of protein was prepared and western blotting was conducted according to the NuPAGE electrophoresis system (Thermo Fisher Scientific). We used the following primary antibodies: mouse monoclonal anti-dystrophin (MANDYS8; Sigma Aldrich; 1:5000), mouse monoclonal anti-vinculin (V284; Millipore; 1:2500) and mouse monoclonal anti-β-actin (sc-47778; Santa Cruz Biotechnology; 1:10,000).

### 14. Statistical analysis

GraphPad Prism version 7 was used to perofrm Student’s t-tests for all statistical analyses.

## Acknowledgments

We acknowledge the members of the Cohn, Ivakine and Delgado laboratories for their input and technical support. We also would like to thank The Center for Phenogenomics for mouse husbandry and Shagana Visuvanathan for her help with colony management.

## Author Contributions

Conceptualization: TWYW, LP, EAI, RDC. Methodology: LP, TWYW, AA, EM, EH, OS. Formal analysis: TWYW, LP, EM, OS, AA, PDO. Investigation: TWYW, LP, AA, EM, OS. Writing – original draft: LP. Writing – review and editing: LP, EM, OS, PDO, EAI, RDC. Visualization: LP, TWYW, EM, OS. Supervision: EAI, RDC. Project administration: EAI, RDC. Funding acquisition: TWYW, EAI, RDC.

## Data availability

All original data are available from the authors without any restrictions.

